# CK-666 and CK-869 differentially inhibit Arp2/3 iso-complexes

**DOI:** 10.1101/2023.11.26.568719

**Authors:** LuYan Cao, Shaina Huang, Angika Basant, Miroslav Mladenov, Michael Way

## Abstract

The inhibitors, CK-666 and CK-869, are widely used to probe the function of actin nucleation by the Arp2/3 complex *in vitro* and in cells. However, in mammals, the Arp2/3 complex consists of 8 iso-complexes, as three of its subunits (Arp3, ArpC1, ArpC5) are encoded by two different genes. Here, we used recombinant Arp2/3 with defined composition to assess the activity of CK-666 and CK-869 against iso-complexes. We demonstrate that both inhibitors prevent linear actin filament formation when ArpC1A- or ArpC1B-containing complexes are activated by SPIN90. In contrast, inhibition of actin branching depends on iso-complex composition. Both drugs prevent actin branch formation by complexes containing ArpC1A, but only CK-869 can inhibit ArpC1B-containing complexes. Consistent with this, in bone marrow-derived macrophages which express low levels of ArpC1A, CK-869 but not CK-666, impacted phagocytosis and cell migration. CK-869 is also only able to inhibit Arp3-but not Arp3B-containing iso-complexes. Our findings have important implications for the interpretation of results using CK-666 and CK-869, given that the relative expression levels of ArpC1 and Arp3 isoforms in cells and tissues remains largely unknown.

## Introduction

The Arp2/3 complex is an important evolutionarily conserved nucleator of actin filaments (Machesky *et al*, 1997; Papalazarou & Machesky, 2021; Welch *et al*, 1997). The complex regulates the architecture and dynamics of actin cytoskeleton by initiating the formation of both branched and linear actin filaments (Cao & Way, 2024; Espinoza-Sanchez *et al*, 2018; Gautreau *et al*, 2022; Mullins *et al*, 1998; Wagner *et al*, 2013). When activated by class I nucleating promoting factors (NPFs), such as WAVE, N-WASP and WASH, the Arp2/3 complex binds to the side of a pre-existing (mother) actin filament and nucleates the formation of a new (daughter) actin branch (Espinoza-Sanchez *et al*., 2018; Molinie & Gautreau, 2018; Smith *et al*, 2013). Alternatively, Arp2/3 activation by SPIN90 (also known as WISH, NCKIPSD, Dip1) inhibits the association of the complex with an existing actin filament and generates a new linear actin filament (Luan *et al*, 2018; Wagner *et al*., 2013). Through these two activities, the Arp2/3 complex plays essential roles in various cell processes, including endocytosis, cell migration, regulation of the actin cortex and DNA repair (Cao *et al*, 2020; Hinze & Boucrot, 2018; Papalazarou & Machesky, 2021; Schrank *et al*, 2018; Wu *et al*, 2012).

The Arp2/3 complex consists of seven subunits, including two <u>A</u>ctin <u>R</u>elated <u>P</u>roteins: Arp2, Arp3 and ArpC1-ArpC5. Upon activation by class I NPFs, the ArpC1-5 subunits interact with the mother actin filament while Arp2 and Arp3 act as a template to nucleate a new daughter filament (Chou *et al*, 2022; Ding *et al*, 2022; Fassler *et al*, 2020). In mammals but not lower eukaryotes, such as yeast and amoeba, Arp3, ArpC1 and ArpC5 exist as two different isoforms that are encoded by separate genes (Balasubramanian *et al*, 1996; Jay *et al*, 2000; Millard *et al*, 2003). In humans, Arp3 and Arp3B, ArpC1A and ArpC1B, ArpC5 and ArpC5L are 91, 67 and 67% identical respectively (Abella *et al*, 2016). Hence, in mammals, the Arp2/3 complex is not a single species but a group of eight different iso-complexes. Moreover, these iso-complexes have different molecular properties as Arp2/3 complexes containing ArpC1B and ArpC5L are significantly better at stimulating actin assembly than those with ArpC1A and ArpC5 (Abella *et al*., 2016), while actin networks assembled by Arp3B complexes disassemble faster than those formed by Arp3 complexes (Galloni *et al*, 2021). Structural analysis reveals that the higher branching efficiency of ArpC5L complexes is mediated by its disordered N-terminus (von Loeffelholz *et al*, 2020). Not surprisingly given their different properties, Arp2/3 iso-complexes have different cellular functions (Fassler *et al*, 2023; Molinie *et al*, 2019; Roman *et al*, 2017; Sadhu *et al*, 2023). The importance of Arp2/3 iso-complexes in cell and tissue homeostasis is also underscored by the observation that loss of function mutations in human ArpC1B lead to severe inflammation and immunodeficiency (Brigida *et al*, 2018; Kahr *et al*, 2017; Kuijpers *et al*, 2017; Randzavola *et al*, 2019; Somech *et al*, 2017; Volpi *et al*, 2019). It has also recently been reported that loss of ArpC5 expression results in multiple congenital anomalies, recurrent infections, systemic inflammation, and mortality (Nunes-Santos *et al*, 2023; Sindram *et al*, 2023).

CK-666 and CK-869 are widely used as chemical inhibitors to probe the function of Arp2/3 complex in biochemical assays, cells and tissues (Nolen *et al*, 2009; Rotty *et al*, 2013; Yang *et al*, 2012). CK-666 binds between Arp2 and Arp3 to block the movement of these two subunits, which is required for activation of the Arp2/3 complex (Baggett *et al*, 2012; Hetrick *et al*, 2013; Rouiller *et al*, 2008). CK-869 inhibits this activating conformational change by binding to a hydrophobic pocket in Arp3 (Hetrick *et al*., 2013). CK-869 has also been reported to inhibit microtubule assembly *in vitro* and cells (Yamagishi *et al*, 2018). Taking their different properties into consideration, we wondered whether all Arp2/3 iso-complexes can be inhibited by CK-666 and CK-869. In addition, it also remains to be established whether both drugs are equally effective against class 1 NPF- and SPIN90-mediated activation of Arp2/3 iso-complexes. To address these questions, we performed *in vitro* pyrene and TIRF assays, to examine the impact of CK-666 and CK-869 on the ability of defined recombinant Arp2/3 iso-complexes to nucleate actin when activated by the VCA domain of N-WASP or SPIN90. We found that the ability of CK-666 and CK-869 to inhibit Arp2/3 iso-complexes depends on their subunit composition and activation mechanism. Our observations have important implications for the interpretation of results from experiments using CK666 and CK869 to inhibit mammalian Arp2/3 complex both in cells and using purified complexes with unknown isoform composition. They also provide insights into the differences of Arp2/3 iso-complex activation by VCA domains and SPIN90.

## Results and discussion

### CK-869, but not CK-666, fully inhibits Vaccinia-induced actin polymerisation

During their egress from infected cells, newly assembled vaccinia virus virions that have fused with the plasma membrane and remain attached to the cell induce actin polymerization to enhance their spread into neighbouring uninfected cells (Cudmore *et al*, 1995; Cudmore *et al*, 1996; Frischknecht *et al*, 1999; Leite & Way, 2015). These virions stimulate actin assembly by locally activating Src and Abl family kinases, which results in the recruitment of a signalling network involving the class I NPF, N-WASP that activates Arp2/3 complex dependent actin polymerization (Donnelly *et al*, 2013; Frischknecht *et al*., 1999; Humphries *et al*, 2014; Newsome *et al*, 2004; Newsome *et al*, 2006; Reeves *et al*, 2005; Scaplehorn *et al*, 2002). The resulting actin polymerisation beneath the virion appears as an actin tail when labelled with phalloidin or the branched actin marker cortactin (Figure 1A). Vaccinia induced actin tails have been used as a model system to study actin assembly in the cells (Basant & Way, 2022, 2023; Weisswange *et al*, 2009). We also previously took advantage of vaccinia to demonstrate that Arp2/3 iso-complexes have different cellular activities (Abella *et al*., 2016; Galloni *et al*., 2021). We decided to use the same model to examine whether CK-666 and CK-869 differentially impact the ability of the vaccinia to promote branched actin network assembly. Examination of the literature reveals that these drugs are typically used at 100 µM in cell-based assays. Given this, we treated vaccinia infected HeLa cells 8 hours post infection with 50, 100, 200 and 300 µM CK-666 or CK-869 for 60 minutes and quantified the ability of the virus to stimulate branched actin assembly using cortactin as a marker (Figure 1B, Supplementary Figure 1). We found that while increasing doses of CK-869 result in a near-complete loss of virus induced actin assembly, CK-666 decreased (∼30%) but did not abolish actin polymerisation even at 300 µM (Figure 1B, C). The actin tails induced by the virus in the presence of CK-666, however, were shorter than in the DMSO control (Figure 1A, B). The ability of vaccinia to stimulate actin polymerization depends on their microtubule transport to the cell periphery and fusion with the plasma membrane (Hollinshead *et al*, 2001; Leite & Way, 2015; Newsome *et al*., 2004; Rietdorf *et al*, 2001). The inhibitory effect of CK-869 on actin assembly, however, was not the consequence of the lack of the virus reaching the plasma membrane as extracellular virions were readily detected on the outside of the cell (Figure 1B). Our observations on vaccinia infected HeLa cells demonstrate that CK-666 and CK-869 do not have the same impact on Arp2/3 driven actin assembly.

**Figure 1.**
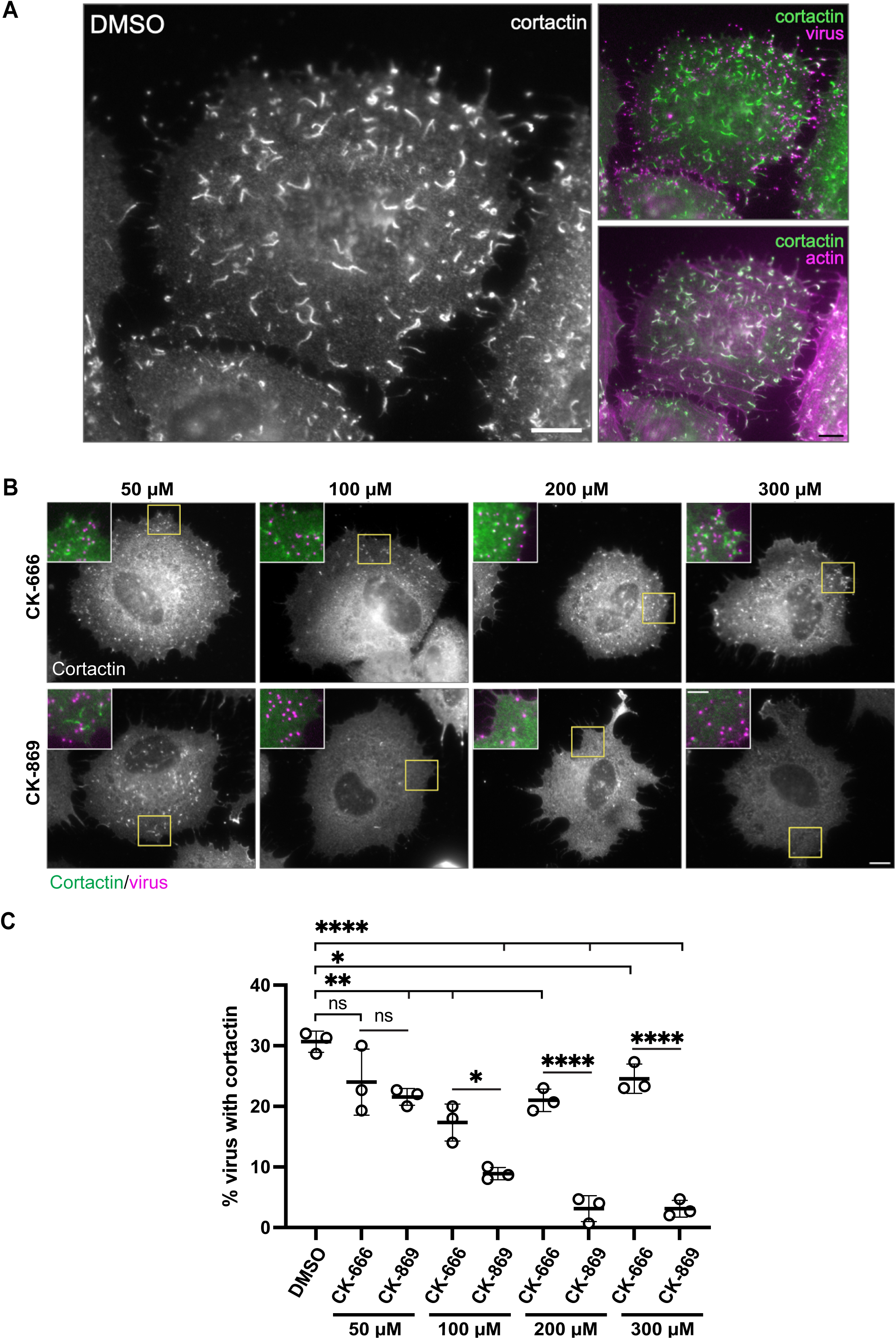
CK-869, but not CK-666 fully inhibits vaccinia-induced actin polymerisation. **A**. Representative immunofluorescence image of a HeLa cell infected with vaccinia virus at 9 hours post-infection and labelled with an antibody against cortactin and stained with phalloidin to visualize F-actin. Extra-cellular virus attached to plasma membrane are labelled with an antibody against the viral protein B5. Scale bar = 10 μm. **B**. Representative images showing localization of cortactin in HeLa cells infected with vaccinia at 9 hours post-infection after 1 hour treatment with CK-666 or CK-869 at the specified concentrations. Scale bar = 10 μm. The inset colour images show 2x magnifications of the yellow boxed regions with cortactin (green) and extra-cellular virus (magenta). **C**. The graph shows quantification of the mean number of extracellular virus co-localising with cortactin from three independent repeats with the error bar representing standard deviation. 10 cells and 150 virus particles were analysed in each experiment. A two-tailed unpaired t-test was used to analyse the statistical significance between different drugs at the same concentration as well as between the drug and DMSO control. ns = not significant. * = p-value < 0.05. ** = p-value < 0.01. *** = p-value < 0.001. **** = p-value < 0.0001.

### CK-666 does not inhibit ArpC1B-containing Arp2/3 complexes

It is not straightforward to deconvolve results from vaccinia infected cells as the precise levels of the 8 different Arp2/3 iso-complexes, which may not be equally inhibited by CK-666 or CK-869 are not known in HeLa cells. We therefore examined the impact of the two inhibitors on defined recombinant Arp2/3 complexes activated by GST-N-WASP-VCA using *in vitro* pyrene actin polymerisation assays (Figure 2, Supplementary Figures 2 and 3). We initially used Arp2/3 iso-complexes containing Arp3 given it is much more abundant than Arp3B in cells and tissues (Galloni *et al*., 2021; Hein *et al*, 2015; Kulak *et al*, 2014). We found both inhibitors suppress activation of ArpC1A containing Arp2/3 complexes but only CK-869 and not CK-666 can inhibit those with ArpC1B (Figure 2A). A similar difference in inhibition was seen in complexes with ArpC5 or ArpC5L (Supplementary Figure 3). The lack of inhibition of ArpC1B complexes by CK-666 was unexpected given that the drug binds between Arp2 and Arp3 to block their movement during Arp2/3 complex activation (Baggett *et al*., 2012; Hetrick *et al*., 2013).

**Figure 2.**
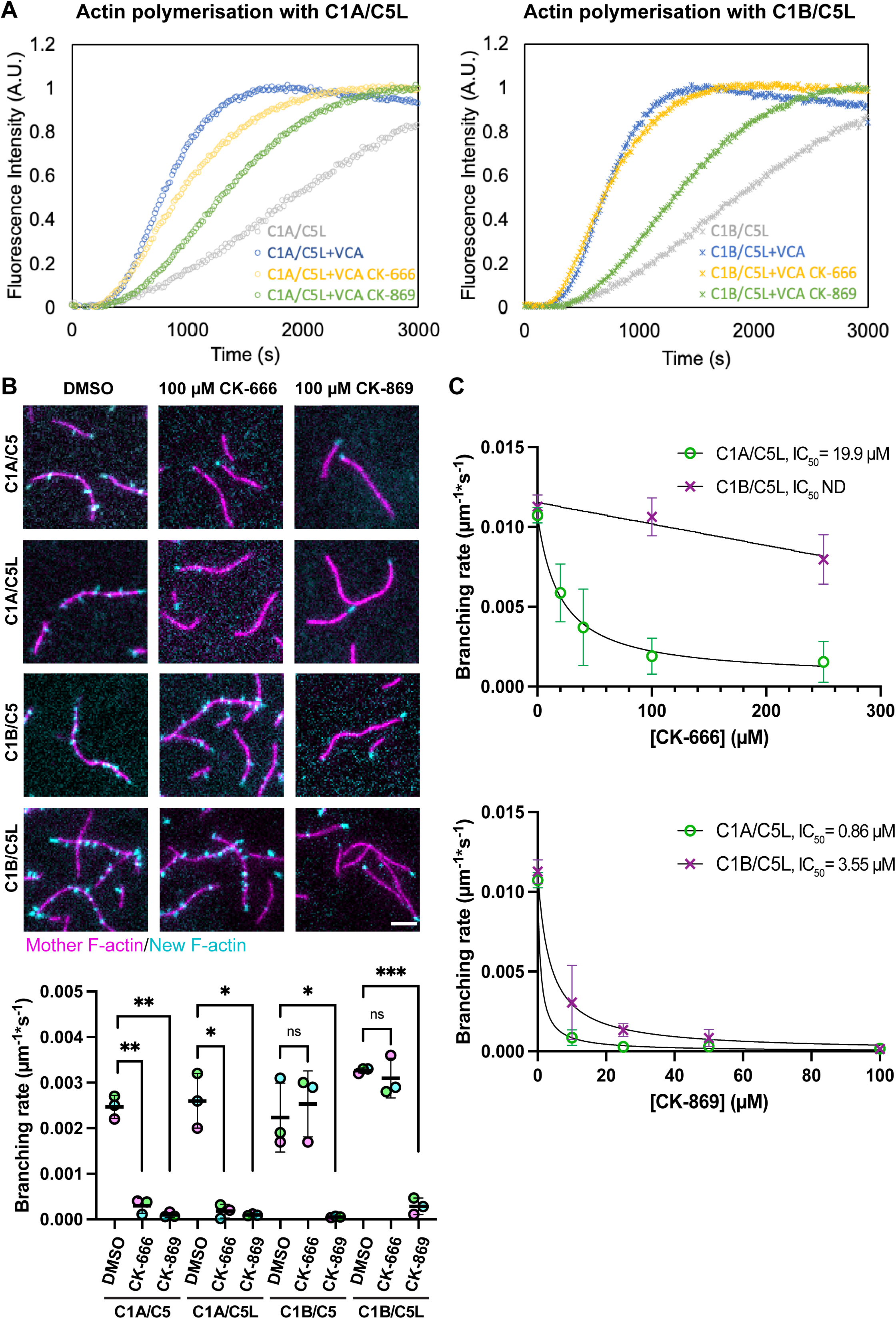
CK-666 does not inhibit ArpC1B containing Arp2/3 complexes. **A**. Representative plots showing the polymerization of pyrene labelled actin (Fluorescence Intensity) when Arp2/3 iso-complexes containing ArpC1A/C5L (left) or ArpC1B/C5L (right) are activated by GST-N-WASP-VCA in the absence (blue) or presence of 100 µM CK-666 (yellow) or CK-869 (green). **B.** Representative TIRF images showing mother filaments (magenta) and new daughter branches (cyan) 2 minutes after Arp2/3 iso-complexes containing ArpC1A/C5, ArpC1A/C5L, ArpC1B/C5 or ArpC1B/C5L were activated by GST-N-WASP-VCA in the absence (DMSO) or presence of 100 µM CK-666 or CK-869. Scale bar = 5 μm. The graph shows the quantification of the mean branching rate for each condition from three independent experiments with the error bar representing standard deviation. Two tailed paired t-test has been applied to analyse the statistical significance. * = p-value < 0.05. *** = p-value < 0.001. ns = not significant. **C.** The branching rate of Arp2/3 complex containing ArpC1A/C5L and ArpC1B/C5L were measured at the indicated concentrations of CK-666 (top) and CK-869 (bottom). The data was fitted with equation Y=Bottom + (Top-Bottom)/(1+(X/IC_50_)) to calculate the half-maximal inhibitory concentration of the drugs (IC_50_), which is indicated (ND = not determined). The points and error bars represent the mean and the standard deviation from at least two independent measurements.

*In vitro* pyrene actin polymerisation assays are an indirect measurement of bulk actin assembly so provide no information at the level of individual filaments. To directly quantify actin branch initiation efficiency, we therefore performed TIRF assays where the generation of new branches is observed over time by incubating fluorescent actin filaments with G-actin labelled with a different fluorophore (Risca *et al*, 2012). We found that 100 µM CK-869 dramatically reduces the branching rate of ArpC1A/C5, ArpC1A/C5L, ArpC1B/C5 and ArpC1B/C5L containing complexes (Figure 2B). In contrast, 100 µM CK-666 was only able to inhibit ArpC1A containing Arp2/3 complexes.

Previous *in vitro* analysis determined that the half-maximal inhibitory concentration value (IC_50_) of CK-666 for human and bovine Arp2/3 purified from platelets and thymus was 4 and 17 µM respectively, while it was 11 µM for CK-869 on bovine Arp2/3 (Nolen *et al*., 2009). As these Arp2/3 complexes are purified from natural sources they are likely to contain different iso-complexes. Given this and our in vitro observations, we determined the IC_50_ for CK-666 and CK-869 on defined Arp2/3 iso-complexes activated by VCA (Figure 2C). We were unable to determine an IC_50_ for ArpC1B/C5L but obtained a value of 19.9 µM for CK-666 with ArpC1A/C5L containing complexes. The lower IC_50_ values we obtained for CK-869 clearly demonstrate that the drug is a significantly better inhibitor than CK-666. Moreover, in an ArpC5L background, CK-869 is more effective against complexes containing ArpC1A (IC_50_ 0.86 µM) than those with ArpC1B (IC_50_ 3.55 µM) (Figure 2C). Although the ArpC5/ArpC5L isoforms are not differentially impacted by either CK-666 or CK-869 (Figure 2B), the latter appears slightly more effective against ArpC5 containing complexes (Supplementary Figure 3B). Our results clearly demonstrate that CK-666 inhibits ArpC1A but not ArpC1B containing Arp2/3 complexes from generating actin branches, whereas CK-869 inhibits both ArpC1 iso-complexes. The results from our *in vitro* assays offer an explanation why CK-666 does not fully suppress vaccinia induced actin polymerization in HeLa cells, which express ∼ 2.3 times as much ArpC1B as ArpC1A (Abella *et al*., 2016).

### CK-666 has no impact on macrophage phagocytosis and motility

Single-cell transcriptomics, tissue specific mRNA expression and proteomic analysis indicate that ArpC1B is highly expressed in immune cells compared to ArpC1A (Kahr *et al*., 2017; Karlsson *et al*, 2021; Kim *et al*, 2014; Wu *et al*, 2009). Given this, we predicted that in contrast to CK-869, CK-666 will have little or no effect on Arp2/3 complex dependent activities in immune cells. After incubating murine bone marrow-derived macrophages with 100 µM of Arp2/3 inhibitor, we observed that the morphology of macrophages treated with CK-869 changed significantly, with cells rounding up to become less spread (Figure 3A, B, Movie 1). In contrast, CK-666 treated macrophages were indistinguishable from DMSO treated controls. In addition, treatment with CK-666 had no impact on macrophage motility (Figure 3C, Movie 1). CK-869, on the other hand, unexpectedly increases macrophage movement. However, a similar increase in motility has been observed in macrophages that contain no functional Arp2/3 complex because they lack ArpC2 (Rotty *et al*, 2017). This study demonstrated that this effect was dependent on myosin II and suggested that the increased motility was due to weakened cell adhesion and enhanced cell contractility. Rotty et al also found that the FcR phagocytic capability of CK-666 treated macrophages was similar to DMSO-treated controls. Extending this observation, we found that CK-869 but not CK-666 decreased the ability of macrophages to phagocytose zymosan compared to DMSO-treated controls (Figure 3D, Supplementary Figure 4). Our observations in macrophages suggest that the ability of CK-666 to inhibit Arp2/3 dependent processes will depend on the relative expression of ArpC1A compared to ArpC1B, which will vary across cell types and tissues.

**Figure 3.**
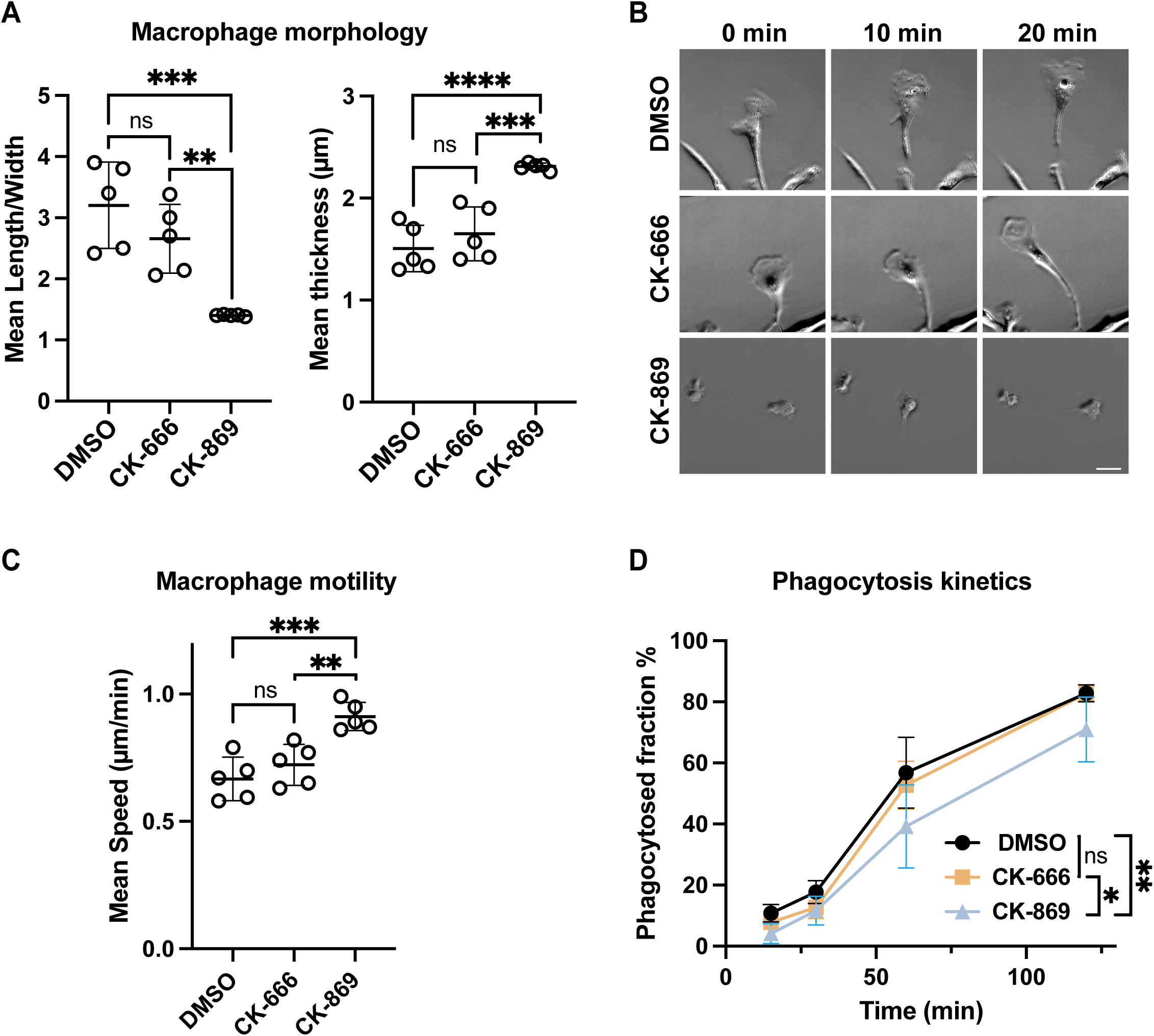
CK666 has no adverse impact on murine bone marrow-derived macrophages. **A.** Treatment of murine bone marrow-derived macrophages (BMDM) with 100 µM CK-869 but not DMSO or CK-666 induces cell rounding. The graphs show the mean ratio of length and width as well as cell thickness from five independent experiments with error bars representing the standard deviation. In each experimental condition, at least 30 cells were randomly chosen and analysed. Two tailed unpaired t-test was used to analyse the statistical significance. **B.** Representative phase images of BMDM treated with DMSO, CK-666 or CK-869 over 20 minutes. Scale bar = 20 µm. **C.** Quantification of the mean migration speed of BMDM treated with DMSO, CK-666 or CK-869 from five independent experiments with error bars representing the standard deviation. In each experimental condition, at least 30 cells were randomly chosen and analysed. Two tailed unpaired t-test was used to analyse the statistical significance. **D**. The graph shows the quantification of the phagocytosis efficiency of zymosan by murine bone marrow-derived macrophages treated with DMSO, CK-666 or CK-869 over 120 minutes. In each experimental condition, 10^4^ cells were analysed using flow cytometry. Each point represents the mean of three independent technical repeats and error bars represent the standard deviation. Two-way ANOVA test has been applied to analyse the statistical significance. ** = p-value < 0.01. *** = p-value < 0.001. **** means p-value < 0.0001. ns = not significant.

### CK-666 and CK-869 do not disrupt the Arp2/3 complex binding to F-actin or VCA

Structural data indicate that neither CK-666 nor CK-869 binds directly to the ArpC1 subunit (Nolen *et al*., 2009). The divergent effects of CK-666 on ArpC1A and ArpC1B within the Arp2/3 complex remains enigmatic. A possible explanation may be that the drugs affect the interaction between Arp2/3 iso-complexes and actin filaments indirectly. To investigate this, we performed actin filament co-sedimentation assays using ArpC1A and ArpC1B-containing complexes in the absence of VCA-mediated activation. We found that regardless of the presence of drugs, comparable quantities of ArpC1A and ArpC1B-containing Arp2/3 complexes co-pelleted with 3 or 15 µM actin filaments (Figure 4A). This indicates that CK-666 and CK-869 do not impair the interaction between the Arp2/3 complex and actin filaments, in the context of different ArpC1 isoforms. However, when assays were performed with 3 µM F-actin, the level of ArpC1B/C5L co-pelleting with actin is a nearly twofold greater than complexes containing ArpC1A/C5L (Figure 4A). This may in part explain why Arp2/3 complex with ArpC1B/C5L is a more efficient branch generator than those with ArpC1A/C5L (Figure 2B) (Abella *et al*., 2016).

**Figure 4.**
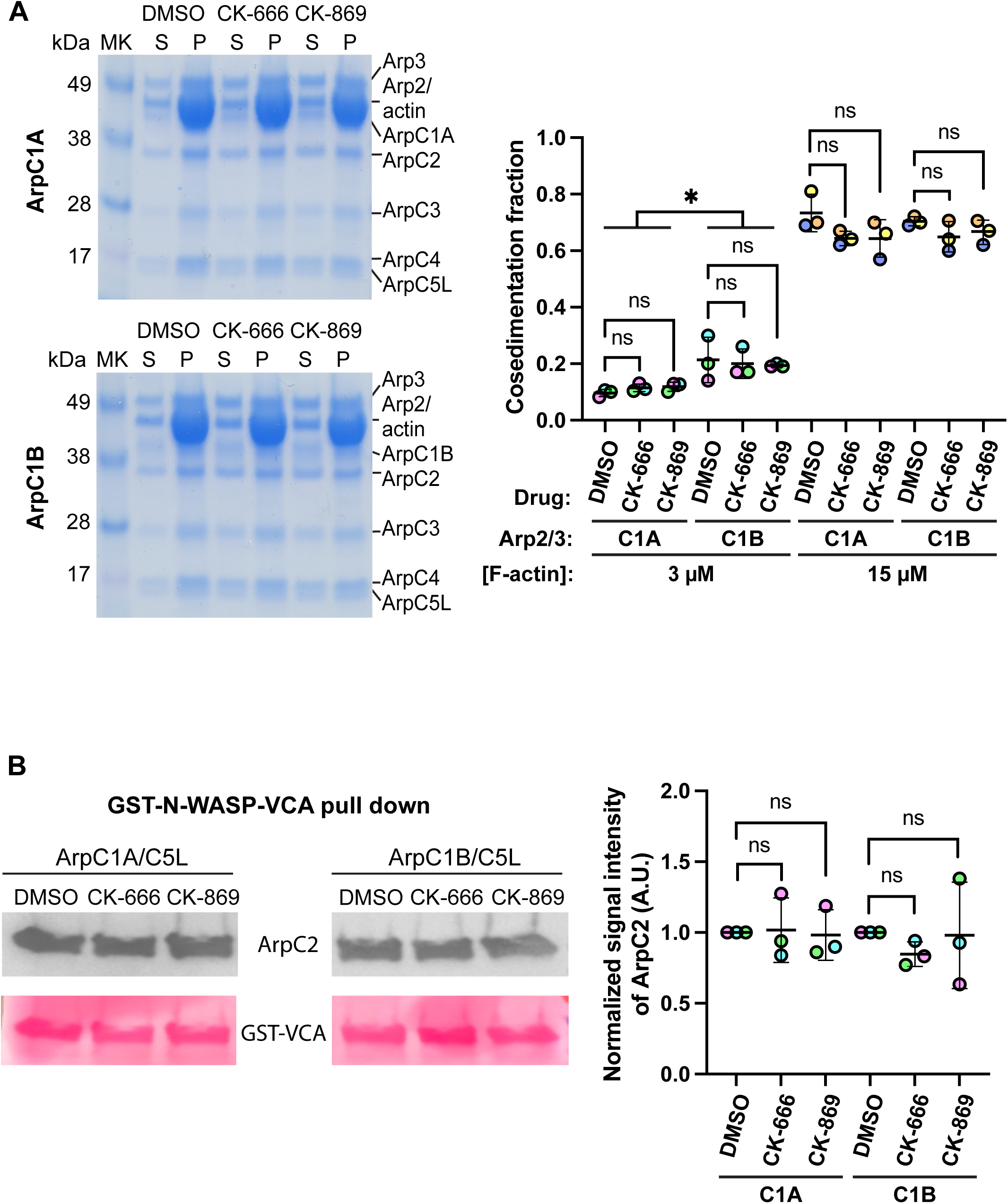
Analysis of Arp2/3 complex binding to actin filaments or VCA. **A.** Left panel shows representative Coomassie stained protein gels of the co-sedimentation of Arp2/3 complexes containing ArpC5L together with ArpC1A (top) or ArpC1B (bottom) with 15 µM actin (P = pellet) in the absence (DMSO) or presence of 100 µM CK-666 or CK-869. The supernatant (S) contains unbound Arp2/3 complexes and non-pelleted G-actin. MK represents the molecular weight markers. Right graph shows the quantification of the mean co-sedimentation fraction of the indicated Arp2/3 complexes in the absence (DMSO) or presence of CK-666 or CK-869 from three independent experiments at 3 and 15 µM actin, with the error bars indicating the standard deviation. Two tailed paired t-test was used to analyse the statistical significance. * = p-value <; 0.05. ns = not significant. **B.** Immunoblots (left panels) using an ArpC2 antibody demonstrate that 100 µM CK-666 or CK-869 does not impact the interaction of Arp2/3 complexes ArpC5L together with ArpC1A (top) or ArpC1B (bottom) with GST-N-WASP-VCA (Ponceau - red). Right graph shows the quantification of the mean pull down fraction of the indicated Arp2/3 complexes in the absence (DMSO) or presence of CK-666 or CK-869 from three independent experiments, with the error bars indicating the standard deviation. Two tailed paired t-test was used to analyse the statistical significance. ns = not significant.

Previous observations indicate that CK-666 and CK-869 do not impact VCA binding (Hetrick *et al*., 2013). However, in this study the Arp2/3 complex will be a mixture of iso-complexes as it was isolated from bovine brain (Abella *et al*., 2016). We therefore performed GST-N-WASP-VCA pulldown assays to capture ArpC1A and ArpC1B containing complexes in the presence or absence of Arp2/3 inhibitors to determine if there are iso-complex specific differences. We found that the presence of either drug did not alter the quantity of ArpC1A- and ArpC1B-containing Arp2/3 complexes bound to GST-N-WASP-VCA (Figure 4B). These results are consistent with previous observations, irrespective of the presence of distinct ArpC1 isoforms (Hetrick *et al*., 2013). It also suggests that inhibited and uninhibited Arp2/3 iso-complexes will compete for VCA binding in cells, which has important implications depending on the relative levels of ArpC1A and ArpC1B and which inhibitor is used. It would also explain why CK-666 inhibited ArpC1A complexes, which represent about a third of the total Arp2/3 in our HeLa cells (Abella *et al*., 2016), decreased virus induced actin polymerization by ∼30% (Figure 1B, C), even though Arp2/3 is in excess of N-WASP (Hein *et al*., 2015; Kulak *et al*., 2014).

### CK-666 inhibits activation of all Arp2/3 iso-complexes by SPIN90

As CK-666 inhibits ArpC1A, but not ArpC1B, containing Arp2/3 complexes from generating actin branches, we investigated whether the same is true when Arp2/3 is activated to form linear actin filaments by SPIN90. Pyrene assays demonstrate that CK-666 and CK-869 inhibit SPIN90-dependent actin polymerisation with a similar IC_50_ irrespective of the ArpC1 isoform (Figure 5A & Supplementary Figure 5). We also observed by TIRF microscopy that both drugs inhibit actin filament formation by ArpC1A and ArpC1B containing Arp2/3 complexes activated by SPIN90, but only CK-869 completely blocked actin assembly (Figure 5B).

**Figure 5.**
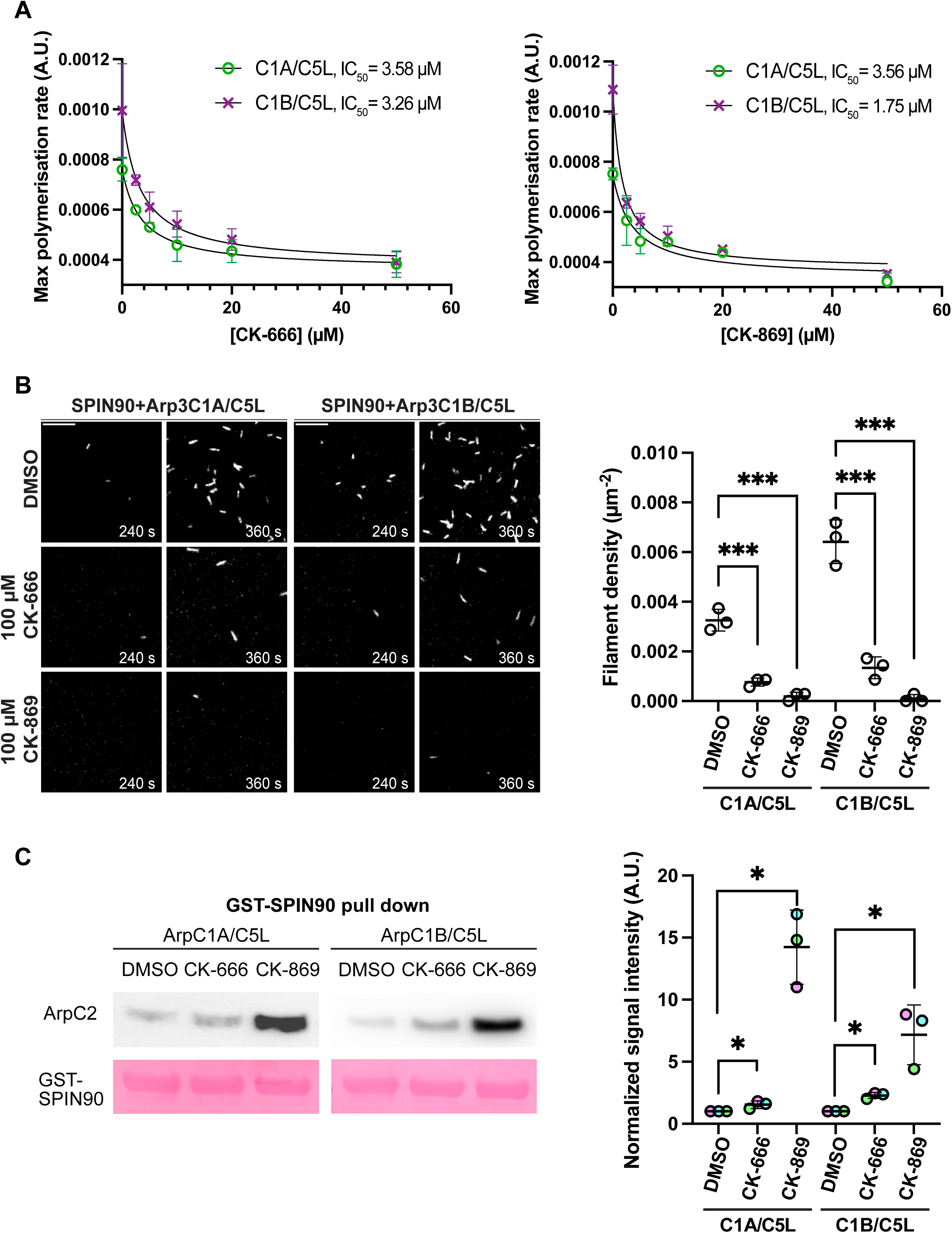
The impact of CK-666 and CK-869 on SPIN90 activation of Arp2/3 complexes. **A.** The maximum rate of pyrene actin polymerisation stimulated by SPIN90-Arp2/3 iso-complexes containing ArpC1A/C5L and ArpC1B/C5L in the presence of the indicated concentrations of CK-666 (top) and CK-869 (bottom). The data was fitted with equation Y=Bottom + (Top-Bottom)/(1+(X/IC50)) to calculate the half-maximal inhibitory concentration of the drugs (IC_50_), which is indicated. The points and error bars represent the mean and the standard deviation of at least two independent measurements. **B.** Representative TIRF images of actin filament assembly induced by Arp2/3 complexes containing ArpC1A/C5L or ArpC1B/C5L after activation by SPIN90-Cter in the absence (DMSO) or presence of 100 µM CK-666 or CK-869 at the indicated times. Scale bar = 10 μm. The graph shows the quantification of the mean filament density at 240 s from three independent experiments with the error bar representing standard deviation. Two tailed unpaired t-test was used to analyse the statistical significance and *** = p-value <; 0.001. **C.** The left panels show immunoblots of GST-SPIN90-Cter (Ponceau - red) pulldown assays of the indicated Arp2/3 iso-complexes (detected by ArpC2 antibody) in the absence (DMSO) or presence of 100 µM CK-666 or CK-869. The right panel shows the quantification of the mean bound Arp2/3 complex normalized to the DMSO control for three independent pulldown assays with the error bar representing standard deviation. Two tailed paired t-test was used to analyse the statistical significance. ns = not significant. * = p-value < 0.5.

Neither drug inhibited the interaction of Arp2/3 with SPIN90 in GST pull down assays (Figure 5C). However, unexpectedly CK-869 promotes the binding of Arp2/3 to SPIN90, with the interaction with ArpC1A and ArpC1B containing complexes increasing ∼15 and 5-fold respectively. There was also a slight increase in binding in the presence of CK-666. A possible explanation is that CK-869 blocks the Arp2/3 complex in an inactive state which favours binding to SPIN90 but prevents further conformational changes required for full activation which would subsequently weaken the association with SPIN90. This would imply that inactive Arp2/3 has a higher affinity for SPIN90 than the active complex as is observed for VCA but not cortactin NtA (Liu *et al*, 2024; Zimmet *et al*, 2020). This may explain why the amount of Arp2/3 pulled down with GST-SPIN90 is reduced in the presence of GST-N-WASP-VCA (Cao *et al*, 2023).

### CK-869 cannot inhibit Arp3B-containing complexes

Our results thus far demonstrate that CK-869 is a better inhibitor of Arp3-containing complexes than CK-666. To explore whether the same is also true for Arp3B containing complexes, we examined the impact of both inhibitors on the ability of Arp3B/ArpC1B/ArpC5L containing Arp2/3 complex to nucleate actin filaments. Quantification of actin assembly in TIRF assays reveals that the branching rates induced by GST-N-WASP-VCA in the presence of CK-666 were indistinguishable from DMSO-treated controls (Figure 6A). Unexpectedly, CK-869 no longer inhibits the Arp3B containing complex, which contrasts with our earlier observations using complexes with Arp3 (Figure 2). Moreover, as observed with Arp3-containing complexes (Figure 5), CK-666 inhibits the ability of SPIN90 to stimulate linear actin filament assembly by Arp3B-containing complexes (Figure 6B). In sharp contrast, Arp3B-containing complexes efficiently generate linear actin filaments in the presence of CK-869. Our *in vitro* data demonstrate that CK-869 cannot inhibit Arp3B-containing complexes, even up to 250 µM in pyrene actin assembly assays (Figure 6C).

**Figure 6.**
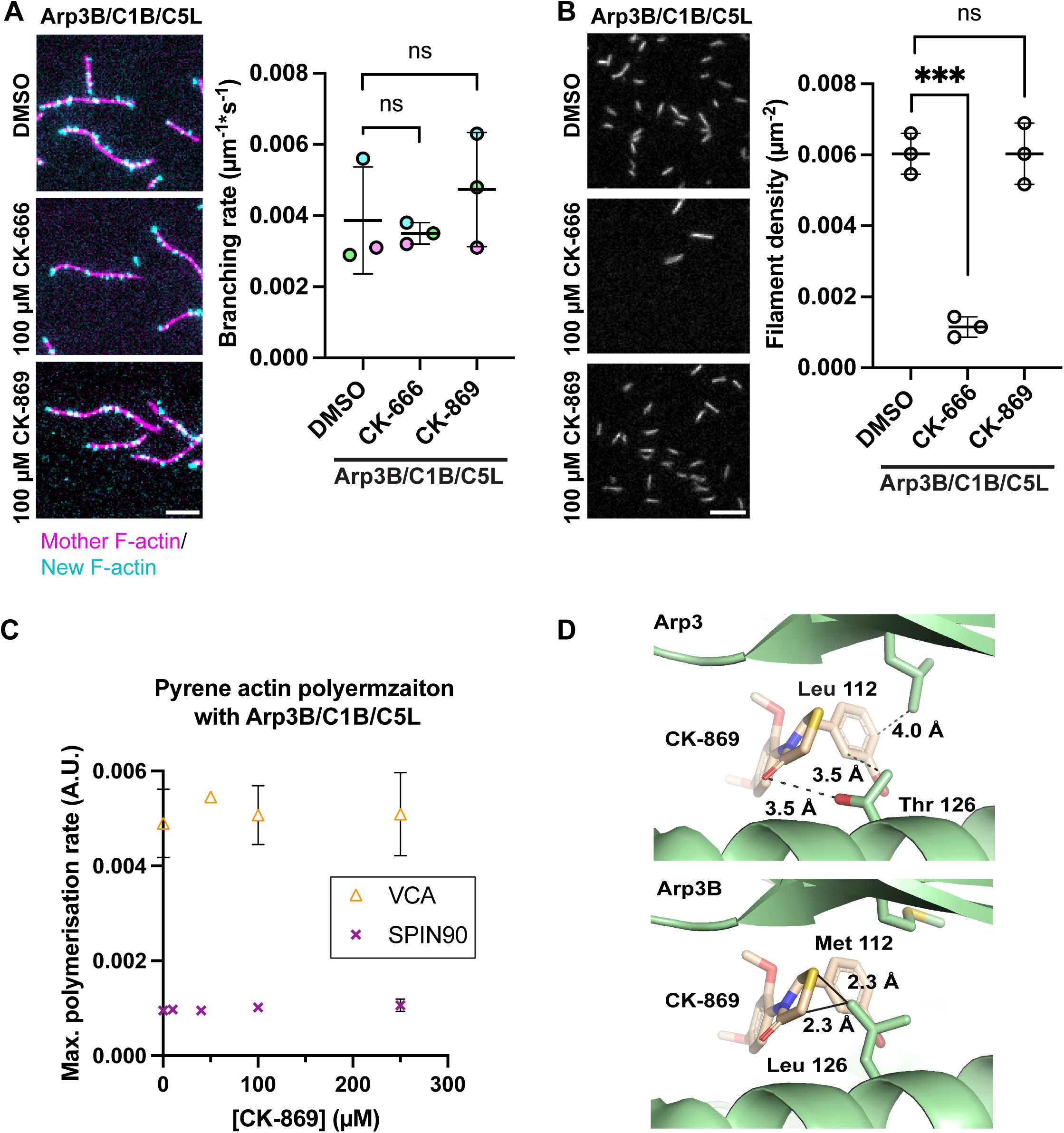
Impacts of drugs on Arp3B containing complexes. **A** Representative TIRF images showing mother filaments (magenta) and newly polymerised daughter actin branches (cyan) 2 minutes after Arp2/3 complex containing Arp3B/ArpC1B/C5L was activated by GST-N-WASP-VCA in the absence (DMSO) or presence of 100 µM CK-666 or CK-869. Scale bar = 5 μm. The graph shows the quantification of the mean branching rate for each condition from three independent experiments with the error bar representing standard deviation. Two tailed paired t-test was used to analyse the statistical significance and ns = not significant. **B.** Representative TIRF images of actin filament assembly induced by Arp2/3 complex containing Arp3B/ArpC1B/C5L after activation by SPIN90-Cter in the absence (DMSO) or presence of 100 µM CK-666 or CK-869 at the indicated times. Scale bar = 10 μm. The graph shows the quantification of the mean filament density at 240 s from three independent experiments with the error bar representing standard deviation. Two tailed unpaired t-test was used to analyse the statistical significance. ns = not significant. *** presents p-value < 0.001. **C**. The maximum rate of pyrene actin polymerisation stimulated by Arp2/3 complex containing Arp3B/ArpC1B/C5L activated by either GST-N-WASP-VCA or SPIN90 Cter at the indicated CK-869 concentration. The points and error bars represent the mean and the standard deviation of at least two independent measurements. **D**. The top image shows that in the Arp2/3 complex (PDB: 3ULE), CK-869 has hydrophobic interactions with Leu 112 and Thr 126 of Arp3. The bottom image shows that in Arp3B, Leu 112 and Thr 126 are replaced by Met 112 and Leu 126, the latter of which is predicted to have clashes with CK-869.

Previous structural analysis reveals that CK-666 interacts mostly with Arp2 but forms two hydrophobic interactions with leucine 117 and threonine 119 of Arp3 (Baggett *et al*., 2012). These two residues are conserved in Arp3B, suggesting that the binding of CK-666 will be similar, which is consistent with our in vitro data. In contrast, several residues in the hydrophobic CK-869 binding pocket of Arp3 differ to those in Arp3B (Hetrick *et al*., 2013) (Figure 6D). Notably, the leucine 126 in Arp3B may have direct clashes with CK-869 and preclude its binding leading to a loss of inhibition. A similar lack of inhibition due to non-conserved residues in the CK-869 binding pocket is also observed in Arp2/3 from budding and fission yeast (Nolen *et al*., 2009).

## Conclusions

Using purified components and *in vitro* assays we have demonstrated that CK-666 can inhibit the formation of SPIN90-induced linear actin filaments by all Arp2/3 iso-complexes. However, CK-666 fails to suppress the initiation of actin branch formation by Arp2/3 complexes containing ArpC1B. In contrast, CK-869 prevents Arp3 but not Arp3B-containing complexes from generating both actin branches and linear actin filaments. Our results provide further evidence that Arp2/3 iso-complexes have different molecular properties. This observation also aligns with our cellular findings, where CK-869 exhibited greater efficacy in inhibiting vaccinia-induced actin assembly compared to CK-666. Examination of the literature reveals that most studies typically treat cells with CK-666 rather than CK-869, presumably because the latter is more toxic as it will result in complete loss of Arp2/3 function.

Our data demonstrate that both drugs have high affinity for Arp2/3 iso-complexes containing Arp3 (Figure 5A). However, CK-666 and CK-869 have a different impact on Arp2/3 iso-complexes, depending on whether they are activated by VCA or SPIN90. Both drugs can effectively inhibit Arp2/3-SPIN90 generating actin filaments, while simultaneously enhancing, to varying degrees, the binding of Arp2/3 complexes to SPIN90. This implies that SPIN90 may induce a relatively weak activation of Arp2/3 complexes, which is more susceptible to inhibition. The observation that CK-666 has no discernible effect on the binding of Arp2/3 to either VCA or mother filaments indicates that CK-666 primarily disrupts the conformational changes of Arp2 and Arp3 to prevent complex activation as suggested by the structural data (Baggett *et al*., 2012). Nevertheless, ArpC1B but not ArpC1A containing complexes are able to adopt an activated conformation to generate actin branches on binding VCA even though they are bound to CK-666. This would suggest that the energy barrier between inactive and active conformations is lower for ArpC1B containing Arp2/3 complexes than those with ArpC1A. It is also possible that the conformation change required for activation displaces CK-666 from Arp2/3 iso-complexes containing ArpC1B. Structural analysis of Arp2/3 iso-complexes in transitional states of complex activation are required to fully understand their different behaviour.

The pivotal role of Arp2/3 complexes in many cellular processes that can become dysregulated in pathological conditions, highlights the ongoing need to screen for additional Arp2/3 inhibitors. Suppressing cancer cell migration and tumour metastasis, two ArpC2 binding inhibitors, were recently identified by revisiting FDA-approved drugs (Choi *et al*, 2019; Yoon *et al*, 2019). Several CK-666 structural analogues have been reported to have inhibitory effect on the Arp2/3 complex and two of them have better *in vivo* efficiency compared with CK-666 (Fokin *et al*, 2022). Given our results, it is possible that these drugs have different inhibitory effects on Arp2/3 iso-complexes. In summary, our results have important implications for the interpretation of experiments with CK-666 and CK-869 but also illuminate the potential utility of these inhibitors in exploring the cellular functions of specific Arp2/3 iso-complexes.

## Materials and methods

### Vaccinia virus infection and drug treatment

HeLa cells plated on fibronectin-coated coverslips were infected with Western Reserve Vaccinia virus in serum-free Minimum Essential Medium (MEM) at multiplicity of infection (MOI) = 2 (Frischknecht *et al*., 1999). After one hour at 37°C, the serum-free MEM was removed and replaced with complete MEM. At 8 hours post-infection, media was replaced with complete MEM containing CK-666 (Sigma, SML0006-5MG) or CK-869 (Sigma, C9124-5MG) or DMSO at desired concentrations for 1 hour prior to fixation.

### Immunofluorescence and actin tail quantification

As previously described (Basant & Way, 2022), cells were fixed with 4% paraformaldehyde in PBS for 10 min, blocked in cytoskeletal buffer (1 mM MES, 15 mM NaCl, 0.5 mM EGTA, 0.5 mM MgCl_2_, and 0.5 mM glucose, pH 6.1) containing 2% (vol/vol) fetal calf serum and 1% (wt/vol) BSA for 30 min. To visualize extra-cellular vaccinia virus particles, prior to permeabilization with detergent, the cells were stained with a monoclonal antibody against B5 (19C2, rat, 1:1000 (Schmelz *et al*, 1994)) followed by an Alexa Fluor 647 anti-rat secondary antibody (Invitrogen; 1:1000 in blocking buffer). Cells were then permeabilized with 0.1% Triton-X/PBS for 3 min. Branched actin was labelled with cortactin monoclonal antibody (Millipore (p80/85); 1:400), followed by Alexa Fluor 488 conjugated secondary antibody (Invitrogen; 1:1000 in blocking buffer). Actin filaments were labelled with Alexa Fluor 568 phalloidin (Invitrogen; 1:500). Coverslips were mounted on glass slides using Mowiol (Sigma) and imaged on a Zeiss Axioplan2 microscope equipped with a 63x/1.4 NA. Plan-Achromat objective and a Photometrics Cool Snap HQ cooled charge-coupled device camera. The microscope was controlled with MetaMorph 7.8.13.0 software. Quantification of branched actin induced by vaccinia virus was performed using two-colour fixed cell images where cortactin and extracellular virus were labelled. Ten cells were analysed per condition in three independent experiments. Fifteen isolated extracellular virus particles were blindly selected in each image and the presence of actin and cortactin at the virus was determined in the corresponding channels.

### Bone marrow-derived macrophage migration and phagocytosis

All animal work was authorized by UK Home Office project license P7E080263 and personal licenses following the approval by the Animal Welfare and Ethical Review Body of The Francis Crick Institute. Murine bone marrow was extracted from tibia and femur of C57BL/6Jax mice and differentiated *in vitro* for 7 days in macrophage conditional medium (RPMI supplemented with 30% L929 conditioned medium, 20% FBS and 1% P/S) to obtain bone marrow-derived macrophages (BMDM).

To examine the effects of different inhibitors on cell morphology and migration, BMDM were seeded in 24-well glass bottomed plates (Cellvis) at 1*10^4^ cells/well in 500 µL of macrophage conditional medium. Cells were imaged and analyzed using the Livecyte system (Phasefocus) every 10 mins for 24 h at 10 x with two fields of view per well. The imaging chamber was maintained at 37°C with 5% CO_2_. Meanwhile, cells were seeded at 2 *10^4^ cells/well in Glass-bottomed 8-well µ-Slides (ibidi) and imaged using an inverted widefield Nikon Ti2 Eclipse long-term time lapse system with an LED illumination system. The imaging chamber was maintained at 37°C with 5% CO_2_. A Nikon 20× Ph2 planar (plan) apochromatic (apo) (0.75 N.A.) air objective was used to acquire images at a single *z*-slice every 10 min for 5 hours with two fields of view per well. Livecyte automatically quantified the motility and the morphology of macrophages by measuring their moving speed as well as their thickness and their length/width ratio. For each condition, more than 30 cells were quantified. The experiments were repeated five times independently.

To assess the effects of different inhibitors on phagocytosis, BMDM were seeded in 96-well V-bottom plates (Corning) at 1*10^5^ cells/well in 100 µL macrophage conditional medium supplemented with different inhibitors or DMSO as indicated. BMDM were incubated with 25 µg/mL pHrodo zymosan bio-particles (Invitrogen) for up to 120 min at 37°C, 5% CO_2_. Cells were then washed with PBS and stained with an antibody cocktail (Live/Dead Fixable Violet Dead Cell Stain (1:500, Invitrogen), CD11b-BV711 (1:100, BioLegend), F4/80-APC (1:100, eBiosciences)) to identify the live BMDM. Samples were then assessed by flow cytometry and the phagocytic fraction indicated the % of (BMDM internalized the zymosan bio-particles) / (Total BMDM).

### Protein purification and preparation

GST-tagged human N-WASP-VCA (392 – 505, UniProt O00401), human SPIN90 C-terminus (267 - 715, UniProt Q9NZQ3-3), were purified as previously reported (Cao *et al*., 2023). Recombinant human Arp2/3 iso-complexes with defined composition (Uniprot P61160, P61158, Q9P1U1, Q92747, O15143, O15144, O15145, P59998, Q9H9F9, Q98PX5) were purified as previously reported (Baldauf *et al*, 2023). Skeletal muscle alpha-actin was purified from rabbit muscle acetone powder following the published protocol (Spudich & Watt, 1971). Actin was fluorescently labelled on the surface lysine 328, using Alexa-488 or Alexa-594 succinimidyl ester (Life Technologies) as previously reported (Romet-Lemonne *et al*, 2018). Pyrenyl-actin was made by labelling actin with N(1-pyrene)-iodoacetamide (Thermo Fisher Scientific) (Cooper *et al*, 1983). To mimic the use of drugs in cells, Arp2/3 iso-complexes were incubated with 100 µM CK-666 or CK-869 (Sigma, Bio-Techne), or equivalent amount of DMSO at room temperature for 1 hour before being used in in vitro assays.

### Pyrene actin polymerisation assay

Actin polymerisation induced by Arp2/3 and NPF I is very fast. To have a good resolution, 2 nM Arp2/3 iso-complexes, 5 nM GST-N-WASP-VCA and 2 µM G-actin (5% pyrene labelled) was mixed and measured in a Safas Xenius fluorimeter at room temperature (von Loeffelholz *et al*., 2020). The negative control experiment was done without GST-N-WASP-VCA. For experiments with SPIN90, the same concentration of Arp2/3 iso-complexes was used together with 100 nM SPIN90-Cter and 2.5 µM G-actin (5% pyrene labelled). The negative control for these experiments lacked SPIN90-Cter. All actin assembly assays above were performed in a buffer containing 5 mM Tris HCl pH 7.0, 50 mM KCl, 0.2 mM ATP, 1 mM MgCl_2_, 1 mM EGTA and 1 mM DTT in addition to 100 µM CK-666 or CK-869 or equivalent amount of DMSO. To quantify IC_50_ values, Arp2/3 iso-complexes were incubated with DMSO or different drug concentrations for 1 hour. Then 2 nM drug treated Arp2/3 iso-complexes were mixed with 10 nM VCA or 400 nM SPIN90-Cter together with 2.5 µM G-actin (5% pyrene labelled) in the presence of the same concentration of drugs. The experimental buffer contains 5 mM Tris HCl pH 7.0, 50 mM KCl, 1 mM MgCl_2_, 0.2 mM EGTA, 0.2 mM ATP, 10 mM DTT, 1 mM DABCO at room temperature. The maximum polymerisation rate of each pyrene curve was measured and plotted versus defined drug concentration.

### TIRF assays

Deep UV treated coverslips were passivated by mPEG silane overnight. The coverslips were rinsed thoroughly with ethanol and water. Before experiments, dried coverslips were stuck to the glass slides with 3 mm-separated parafilm bands to create flow chambers. The experiments were performed in TIRF buffer: 5 mM Tris HCl pH 7.0, 50 mM KCl, 1 mM MgCl_2_, 0.2 mM EGTA, 0.2 mM ATP, 10 mM DTT, 1 mM DABCO, 0.1% BSA and 0.3% Methylcellulose 4000 cP at 25 °C.

Actin branching assays were performed as previously described (Cao *et al*., 2020). To quantify the branching efficiency, 2.5 µM G-actin (15% Alexa-488 labelled) was pre-incubated in experimental buffer at room temperature for 1 hour to form fluorescent actin filaments. During experiment, 25 nM pre-polymerised actin (15% Alexa-488 labelled) was mixed with 25 nM GST-N-WASP-VCA, 0.5 µM profilin, 2 nM of 1 hour inhibitor-treated Arp2/3 in addition to defined concentration of CK-666 or CK-869 (final concentration) or equivalent amount of DMSO. G-actin (0.5 µM, 15 % Alexa-568 labelled) was added to the system at the last moment. Then the mixture was immediately flowed into the chamber for imaging. The moment when G-actin added into the system was recorded as time 0. Branch generation was observed over time. The number of branches generated per µm of actin filaments per minute was quantified as the branching rate. The quantified branching rates versus defined drug concentrations were plotted to determine the IC_50_. To study the actin nucleation by SPIN90, the 25 nM GST-N-WASP-VCA was replaced 200 nM SPIN90-Cter and 20 nM of inhibitor treated Arp2/3 were used as in the same protocol as above.

Actin branching rate in TIRF assay was quantified by counting the number of generated branches per µm length of actin mother filament per second using Fiji software (Schindelin *et al*, 2012). In each condition, the ten longest mother filaments were chosen to analyse. Three independent repeats were performed. Actin nucleation by SPIN90-Arp2/3 complex in TIRF assay was quantified by measuring the filament density of a randomly picked field of view (59 µm x 59 µm) at 240s. Three independent repeats were performed.

### GST pull down assays

Prior to performing pull-down assays, the Glutathione Sepharose^TM^ 4B resin (GE Healthcare) was incubated with 5% BSA for 5 minutes and washed 3 times with GST pull-down buffer containing 50 mM Tris-HCl pH 7.0, 1 mM MgCl_2_, 0.2 mM EGTA, 0.2 mM ATP, 1 mM DTT, 50 mM KCl, and 5% glycerol. Glutathione Sepharose 4B beads were mixed with 50 µL of 5 µM GST-SPIN90-Cter or GST-N-WASP-VCA, 200 nM drug treated Arp2/3 in addition to 100 µM inhibitor or equivalent amount of DMSO. After 1 hour incubation at room temperature, the resin was then washed 3 times with 300 μL of GST pull-down buffer. The proteins bound to the resin were eluted with 50 μL of 50 mM GSH (reduced glutathione). The sample was analysed by SDS-PAGE followed by western blot using Anti-ArpC2 (Sigma) to detect Arp2/3 complexes. Amersham imager AI600 was used to develop the western blot results. The amount of GST-tagged protein eluted from the resin was detected by ponceau. The negative control experiment was done in the same way except 50 μL of 5 μM Glutathione Sepharose^TM^ 4B attached GST was used instead of GST-SPIN90-Cter or GST-N-WASP-VCA.

### F-Actin co-sedimentation assays

G-actin was incubated in F-buffer (5 mM Tris pH 7.0, 100 mM KCl, 0.2 mM ATP, 1 mM MgCl_2_, and 1 mM DTT) at room temperature for 1 hour before 3 µM or 15 µM of the polymerised actin filaments were mixed with 4 µM inhibitor treated Arp2/3 iso-complexes in F-buffer in the presence of 100 µM CK-666 or CK-869 or equivalent amount of DMSO at room temperature for 1 hour. The supernatant and pellet were then separated by ultracentrifuge at 70000 rpm at 25 °C for 30 minutes. Pellet was resuspended with 50 µL of F-buffer and the same amount (20 µL) of supernatant and pellet was loaded on a SDS page. The negative control only has Arp2/3 iso-complexes without F-actin. Fiji software was used to quantify the Coomassie blue gels (Schindelin *et al*, 2012).

### Protein structure analysis

Pymol is used to visualize and analyse the structure of drug bound Arp2/3 complex (Schrodinger, 2010). Potential Van-der-Waals clashes were detected by show bumps plugin.

## Acknowledgements

This project has received funding from the European Research Council (ERC) under the European Union’s Horizon 2020 research and innovation programme (grant agreement No 810207 to Michael Way. Luyan Cao was supported by the European Union’s Horizon 2020 Marie Sklodowka-Curie individual fellowship program (H2020-MSCA-IF-101028239 – MolecularArp) and the ERC Synergy grant (No 810207). MW is additionally supported by the Francis Crick Institute, which receives its core funding from Cancer Research UK (CC2096), the UK Medical Research Council (CC2096), and the Wellcome Trust (CC2096). We thank Carolyn Moores (Birkbeck, University of London), Snezhka Oliferenko & Jeremy Carlton (the Francis Crick Institute, London) and Guillaume Romet-Lemonne & Antoine Jegou (Institut Jacques Monod, Paris) for feedback on the manuscript. For the purpose of Open Access, the authors have applied a CC BY public copyright licence to any Author Accepted Manuscript version arising from this submission.

## Supplementary Figure Legends

**Figure S1. CK-869, but not CK-666, fully inhibits Vaccinia-induced actin polymerisation.** Immunofluorescence images of the actin cytoskeleton (visualised with phalloidin) of HeLa cells infected with Vaccinia virus at 9 hours post-infection after 1-hour incubation with indicated concentrations of CK-666 and CK-869. The images correspond to the actin cytoskeleton of the cortactin images in Figure 1B. Scale bar = 10 μm.

**Figure S2.**

Coomassie stained protein gels of the recombinant proteins and Arp2/3 iso-complexes used in our study.

**Figure S3**

**A.** Representative plots showing the polymerization of pyrene actin (Fluorescence Intensity) when Arp2/3 complexes containing ArpC1A/C5 (left) or ArpC1B/C5 (right) are activated by GST-N-WASP in the absence (blue) or presence of 100 µM CK-666 (yellow) or CK-869 (green).

**B.** The branching rate of Arp2/3 complex containing ArpC1B/C5 and ArpC1B/C5L (same as show in Figure 2C) were measured at the indicated CK-869 concentration. The data was fitted with equation Y=Bottom + (Top-Bottom)/(1+(X/IC_50_)) to calculate the half-maximal inhibitory concentration (IC_50_) of the CK-869. The points and error bars represent the mean and the standard deviation of at least two independent measurements.

**Figure S4.**

Representative FACS plots of murine bone marrow-derived macrophage phagocytosis after treatment with DMSO (control) or 100 µM CK-666 or CK-869.

**Figure S5.**

Representative plots showing the polymerization of pyrene actin (Fluorescence Intensity) when Arp2/3 complexes containing ArpC1A/C5L (left) or ArpC1B/C5L (right) are activated by SPIN90-Cter in the absence (blue) or presence of 100 µM CK-666 (yellow) or CK-869 (green).

**Supplementary Movie 1**

Macrophages expressing LifeAct-GFP were imaged after treated with either DMSO or 100 µM CK-666 or CK-869 for 1 hour. Images were taken every 10 minutes for 5 hours. Scale bar = 100 µm.

## References

Abella JV, Galloni C, Pernier J, Barry DJ, Kjaer S, Carlier MF, Way M (2016) Isoform diversity in the Arp2/3 complex determines actin filament dynamics. Nat Cell Biol 18: 76–86

Baggett AW, Cournia Z, Han MS, Patargias G, Glass AC, Liu SY, Nolen BJ (2012) Structural characterization and computer-aided optimization of a small-molecule inhibitor of the Arp2/3 complex, a key regulator of the actin cytoskeleton. ChemMedChem 7: 1286–1294

Balasubramanian MK, Feoktistova A, McCollum D, Gould KL (1996) Fission yeast Sop2p: a novel and evolutionarily conserved protein that interacts with Arp3p and modulates profilin function. EMBO J 15: 6426–6437

Baldauf L, Frey F, Arribas Perez M, Idema T, Koenderink GH (2023) Branched actin cortices reconstituted in vesicles sense membrane curvature. Biophys J 122: 2311–2324

Basant A, Way M (2022) The relative binding position of Nck and Grb2 adaptors impacts actin-based motility of Vaccinia virus. Elife 11: e74655

Basant A, Way M (2023) The amount of Nck rather than N-WASP correlates with the rate of actin-based motility of Vaccinia virus. Microbiol Spectr 11: e0152923

Brigida I, Zoccolillo M, Cicalese MP, Pfajfer L, Barzaghi F, Scala S, Oleaga-Quintas C, Alvarez-Alvarez JA, Sereni L, Giannelli S et al (2018) T-cell defects in patients with ARPC1B germline mutations account for combined immunodeficiency. Blood 132: 2362–2374

Cao L, Ghasemi F, Way M, Jegou A, Romet-Lemonne G (2023) Regulation of branched versus linear Arp2/3-generated actin filaments. EMBO J 42: e113008

Cao L, Way M (2024) The stabilization of Arp2/3 complex generated actin filaments. Biochem Soc Trans 52: 343–352

Cao L, Yonis A, Vaghela M, Barriga EH, Chugh P, Smith MB, Maufront J, Lavoie G, Meant A, Ferber E et al (2020) SPIN90 associates with mDia1 and the Arp2/3 complex to regulate cortical actin organization. Nat Cell Biol 22: 803–814

Choi J, Lee YJ, Yoon YJ, Kim CH, Park SJ, Kim SY, Doo Kim N, Cho Han D, Kwon BM (2019) Pimozide suppresses cancer cell migration and tumor metastasis through binding to ARPC2, a subunit of the Arp2/3 complex. Cancer Sci 110: 3788–3801

Chou SZ, Chatterjee M, Pollard TD (2022) Mechanism of actin filament branch formation by Arp2/3 complex revealed by a high-resolution cryo-EM structureof the branch junction. Proc Natl Acad Sci U S A 119: e2206722119

Cooper JA, Walker SB, Pollard TD (1983) Pyrene actin: documentation of the validity of a sensitive assay for actin polymerization. J Muscle Res Cell MoAl 4: 253–262

Cudmore S, Cossart P, Griffiths G, Way M (1995) Actin-based motility of vaccinia virus. Nature 378: 636–638

Cudmore S, Reckmann I, Griffiths G, Way M (1996) Vaccinia virus: a model system for actin-membrane interactions. J Cell Sci 109: 1739–1747

Ding B, Narvaez-Ortiz HY, Singh Y, Hocky GM, Chowdhury S, Nolen BJ (2022) Structure of Arp2/3 complex at a branched actin filament junction resolved by single-particle cryo-electron microscopy. Proc Natl Acad Sci U S A 119: e2202723119

Donnelly SK, Weisswange I, Zettl M, Way M (2013) WIP provides an essential link between Nck and N-WASP during Arp2/3-dependent actin polymerization. Curr Biol 23: 999–1006

Espinoza-Sanchez S, Metskas LA, Chou SZ, Rhoades E, Pollard TD (2018) Conformational changes in Arp2/3 complex induced by ATP, WASp-VCA, and actin filaments. Proc Natl Acad Sci U S A 115: E8642–E8651

Fassler F, Dimchev G, Hodirnau VV, Wan W, Schur FKM (2020) Cryo-electron tomography structure of Arp2/3 complex in cells reveals new insights into the branch junction. Nat Commun 11: 6437

Fassler F, Javoor MG, Datler J, Doring H, Hofer FW, Dimchev G, Hodirnau VV, Faix J, Rottner K, Schur FKM (2023) ArpC5 isoforms regulate Arp2/3 complex-dependent protrusion through differential Ena/VASP positioning. Sci Adv 9: eadd6495

Fokin AI, Chuprov-Netochin RN, Malyshev AS, Romero S, Semenova MN, Konyushkin LD, Leonov SV, Semenov VV, Gautreau AM (2022) Synthesis, Screening and Characterization of Novel Potent Arp2/3 Inhibitory Compounds Analogous to CK-666. Front Pharmacol 13: 896994

Frischknecht F, Moreau V, Rottger S, Gonfloni S, Reckmann I, Superti-Furga G, Way M (1999) Actin-based motility of vaccinia virus mimics receptor tyrosine kinase signalling. Nature 401: 926–929

Galloni C, Carra D, Abella JVG, Kjaer S, Singaravelu P, Barry DJ, Kogata N, Guerin C, Blanchoin L, Way M (2021) MICAL2 enhances branched actin network disassembly by oxidizing Arp3B-containing Arp2/3 complexes. J Cell Biol 220: e202102043

Gautreau AM, Fregoso FE, Simanov G, Dominguez R (2022) Nucleation, stabilization, and disassembly of branched actin networks. Trends Cell Biol 32: 421–432

Hein MY, Hubner NC, Poser I, Cox J, Nagaraj N, Toyoda Y, Gak IA, Weisswange I, Mansfeld J, Buchholz F et al (2015) A human interactome in three quantitative dimensions organized by stoichiometries and abundances. Cell 163: 712–723

Hetrick B, Han MS, Helgeson LA, Nolen BJ (2013) Small molecules CK-666 and CK-869 inhibit actin-related protein 2/3 complex by blocking an activating conformational change. Chem Biol 20: 701–712

Hinze C, Boucrot E (2018) Local actin polymerization during endocytic carrier formation. Biochem Soc Trans 46: 565–576

Hollinshead M, Rodger G, Van Eijl H, Law M, Hollinshead R, Vaux DJ, Smith GL (2001) Vaccinia virus utilizes microtubules for movement to the cell surface. J Cell Biol 154: 389–402

Humphries AC, Donnelly SK, Way M (2014) Cdc42 and the Rho GEF intersectin-1 collaborate with Nck to promote N-WASP-dependent actin polymerisation. J Cell Sci 127: 673–685

Jay P, Berge-Lefranc JL, Massacrier A, Roessler E, Wallis D, Muenke M, Gastaldi M, Taviaux S, Cau P, Berta P (2000) ARP3beta, the gene encoding a new human actin-related protein, is alternatively spliced and predominantly expressed in brain neuronal cells. Eur J Biochem 267: 2921–2928

Kahr WH, Pluthero FG, Elkadri A, Warner N, Drobac M, Chen CH, Lo RW, Li L, Li R, Li Q et al (2017) Loss of the Arp2/3 complex component ARPC1B causes platelet abnormalities and predisposes to inflammatory disease. Nat Commun 8: 14816

Karlsson M, Zhang C, Mear L, Zhong W, Digre A, Katona B, Sjostedt E, Butler L, Odeberg J, Dusart P et al (2021) A single-cell type transcriptomics map of human tissues. Sci Adv 7: eabh2169

Kim MS, Pinto SM, Getnet D, Nirujogi RS, Manda SS, Chaerkady R, Madugundu AK, Kelkar DS, Isserlin R, Jain S et al (2014) A drak map of the human proteome. Nature 509: 575–581

Kuijpers TW, Tool ATJ, van der Bijl I, de Boer M, van Houdt M, de Cuyper IM, Roos D, van Alphen F, van Leeuwen K, Cambridge EL et al (2017) Combined immunodeficiency with severe inflammation and allergy caused by ARPC1B deficiency. J Allergy Clin Immunol 140: 273–277 e210

Kulak NA, Pichler G, Paron I, Nagaraj N, Mann M (2014) Minimal, encapsulated proteomic-sample processing applied to copy-number estimation in eukaryotic cells. Nat Methods 11: 319–324

Leite F, Way M (2015) The role of signalling and the cytoskeleton during Vaccinia Virus egress. Virus Res 209: 87–99

Liu T, Cao L, Mladenov M, Jegou A, Way M, Moores CA (2024) Cortactin stabilizes actin branches by bridging activated Arp2/3 to its nucleated actin filament. Nat Struct Mol Biol doi: 10.1038/s41594-023-01205-2. Online ahead of print.

Luan Q, Liu SL, Helgeson LA, Nolen BJ (2018) Structure of the nucleation-promoting factor SPIN90 bound to the actin filament nucleator Arp2/3 complex. EMBO J 37: e100005

Machesky LM, Reeves E, Wientjes F, Mattheyse FJ, Grogan A, Totty NF, Burlingame AL, Hsuan JJ, Segal AW (1997) Mammalian actin-related protein 2/3 complex localizes to regions of lamellipodial protrusion and is composed of evolutionarily conserved proteins. Biochem J 328: 105–112

Millard TH, Behrendt B, Launay S, Futterer K, Machesky LM (2003) Identification and characterisation of a novel human isoform of Arp2/3 complex subunit p16-ARC/ARPC5. Cell MoAl Cytoskeleton 54: 81–90

Molinie N, Gautreau A (2018) The Arp2/3 Regulatory System and Its Deregulation in Cancer. Physiol Rev 98: 215–238

Molinie N, Rubtsova SN, Fokin A, Visweshwaran SP, Rocques N, Polesskaya A, Schnitzler A, Vacher S, Denisov EV, Tashireva LA et al (2019) Cortical branched actin determines cell cycle progression. Cell Res 29: 432–445

Mullins RD, Heuser JA, Pollard TD (1998) The interaction of Arp2/3 complex with actin: nucleation, high affinity pointed end capping, and formation of branching networks of filaments. Proc Natl Acad Sci U S A 95: 6181–6186

Newsome TP, Scaplehorn N, Way M (2004) SRC mediates a switch from microtubule- to actin-based motility of vaccinia virus. Science 306: 124–129

Newsome TP, Weisswange I, Frischknecht F, Way M (2006) Abl collaborates with Src family kinases to stimulate actin-based motility of vaccinia virus. Cell Microbiol 8: 233–241

Nolen BJ, Tomasevic N, Russell A, Pierce DW, Jia Z, McCormick CD, Hartman J, Sakowicz R, Pollard TD (2009) Characterization of two classes of small molecule inhibitors of Arp2/3 complex. Nature 460: 1031–1034

Nunes-Santos CJ, Kuehn H, Boast B, Hwang S, Kuhns DB, Stoddard J, Niemela JE, Fink DL, Pittaluga S, Abu-Asab M et al (2023) Inherited ARPC5 mutations cause an actinopathy impairing cell motility and disrupting cytokine signaling. Nat Commun 14: 3708

Papalazarou V, Machesky LM (2021) The cell pushes back: The Arp2/3 complex is a key orchestrator of cellular responses to environmental forces. Curr Opin Cell Biol 68: 37–44

Randzavola LO, Strege K, Juzans M, Asano Y, Stinchcombe JC, Gawden-Bone CM, Seaman MN, Kuijpers TW, Griffiths GM (2019) Loss of ARPC1B impairs cytotoxic T lymphocyte maintenance and cytolytic activity. J Clin Invest 129: 5600–5614

Reeves PM, Bommarius B, Lebeis S, McNulty S, Christensen J, Swimm A, Chahroudi A, Chavan R, Feinberg MB, Veach D et al (2005) Disabling poxvirus pathogenesis by inhibition of Abl-family tyrosine kinases. Nat Med 11: 731–739

Rietdorf J, Ploubidou A, Reckmann I, Holmstrom A, Frischknecht F, Zettl M, Zimmermann T, Way M (2001) Kinesin-dependent movement on microtubules precedes actin-based motility of vaccinia virus. Nat Cell Biol 3: 992–1000

Risca VI, Wang EB, Chaudhuri O, Chia JJ, Geissler PL, Fletcher DA (2012) Actin filament curvature biases branching direction. Proc Natl Acad Sci U S A 109: 2913–2918

Roman W, Martins JP, Carvalho FA, Voituriez R, Abella JVG, Santos NC, Cadot B, Way M, Gomes ER (2017) Myofibril contraction and crosslinking drive nuclear movement to the periphery of skeletal muscle. Nat Cell Biol 19: 1189–1201

Romet-Lemonne G, Guichard B, Jegou A (2018) Using Microfluidics Single Filament Assay to Study Formin Control of Actin Assembly. Methods Mol Biol 1805: 75–92

Rotty JD, Brighton HE, Craig SL, Asokan SB, Cheng N, Ting JP, Bear JE (2017) Arp2/3 Complex Is Required for Macrophage Integrin Functions but Is Dispensable for FcR Phagocytosis and In Vivo Motility. Dev Cell 42: 498–513.e496

Rotty JD, Wu C, Bear JE (2013) New insights into the regulation and cellular functions of the ARP2/3 complex. Nat Rev Mol Cell Biol 14: 7–12

Rouiller I, Xu XP, Amann KJ, Egile C, Nickell S, Nicastro D, Li R, Pollard TD, Volkmann N, Hanein D (2008) The structural basis of actin filament branching by the Arp2/3 complex. J Cell Biol 180: 887–895

Sadhu L, Tsopoulidis N, Hasanuzzaman M, Laketa V, Way M, Fackler OT (2023) ARPC5 isoforms and their regulation by calcium-calmodulin-N-WASP drive distinct Arp2/3-dependent actin remodeling events in CD4 T cells. Elife 12: e82450

Scaplehorn N, Holmstrom A, Moreau V, Frischknecht F, Reckmann I, Way M (2002) Grb2 and Nck act cooperatively to promote actin-based motility of vaccinia virus. Curr Biol 12: 740–745

Schindelin J, Arganda-Carreras I, Frise E, Kaynig V, Longair M, Pietzsch T, Preibisch S, Rueden C, Saalfeld S, Schmid B et al (2012) Fiji: an open-source planorm for biological-image analysis. Nat Methods 9: 676–682

Schmelz M, Sodeik B, Ericsson M, Wolffe EJ, Shida H, Hiller G, Griffiths G (1994) Assembly of vaccinia virus: the second wrapping cisterna is derived from the trans Golgi network. J Virol 68: 130–147

Schrank BR, Aparicio T, Li Y, Chang W, Chait BT, Gundersen GG, Gottesman ME, Gautier J (2018) Nuclear ARP2/3 drives DNA break clustering for homology-directed repair. Nature 559: 61–66

Schrodinger, LLC, 2010. The PyMOL Molecular Graphics System, Version 1.3r1.

Sindram E, Caballero-Oteyza A, Kogata N, Chor Mei Huang S, Alizadeh Z, Gamez-Diaz L, Fazlollhi MR, Peng X, Grimbacher B, Way M et al (2023) ARPC5 deficiency leads to severe early-onset systemic inflammation and mortality. Dis Model Mech 16: dmm050145

Smith BA, Padrick SB, Doolittle LK, Daugherty-Clarke K, Correa IR, Jr., Xu MQ, Goode BL, Rosen MK, Gelles J (2013) Three-color single molecule imaging shows WASP detachment from Arp2/3 complex triggers actin filament branch formation. Elife 2: e01008

Somech R, Lev A, Lee YN, Simon AJ, Barel O, Schiby G, Avivi C, Barshack I, Rhodes M, Yin J et al (2017) Disruption of Thrombocyte and T Lymphocyte Development by a Mutation in ARPC1B. J Immunol 199: 4036–4045

Spudich JA, WaX S (1971) The regulation of rabbit skeletal muscle contraction. I. Biochemical studies of the interaction of the tropomyosin-troponin complex with actin and the proteolytic fragments of myosin. J Biol Chem 246: 4866–4871

Volpi S, Cicalese MP, Tuijnenburg P, Tool ATJ, Cuadrado E, Abu-Halaweh M, Ahanchian H, Alzyoud R, Akdemir ZC, Barzaghi F et al (2019) A combined immunodeficiency with severe infections, inflammation, and allergy caused by ARPC1B deficiency. J Allergy Clin Immunol 143: 2296–2299

von Loeffelholz O, Purkiss A, Cao L, Kjaer S, Kogata N, Romet-Lemonne G, Way M, Moores CA (2020) Cryo-EM of human Arp2/3 complexes provides structural insights into actin nucleation modulation by ARPC5 isoforms. Biol Open 9: bio054304

Wagner AR, Luan Q, Liu SL, Nolen BJ (2013) Dip1 defines a class of Arp2/3 complex activators that function without preformed actin filaments. Curr Biol 23: 1990–1998

Weisswange I, Newsome TP, Schleich S, Way M (2009) The rate of N-WASP exchange limits the extent of ARP2/3-complex-dependent actin-based motility. Nature 458: 87–91

Welch MD, DePace AH, Verma S, Iwamatsu A, Mitchison TJ (1997) The human Arp2/3 complex is composed of evolutionarily conserved subunits and is localized to cellular regions of dynamic actin filament assembly. J Cell Biol 138: 375–384

Wu C, Asokan SB, Berginski ME, Haynes EM, Sharpless NE, Griffith JD, Gomez SM, Bear JE (2012) Arp2/3 is critical for lamellipodia and response to extracellular matrix cues but is dispensable for chemotaxis. Cell 148: 973–987

Wu C, Orozco C, Boyer J, Leglise M, Goodale J, Batalov S, Hodge CL, Haase J, Janes J, Huss JW, 3rd et al (2009) BioGPS: an extensible and customizable portal for querying and organizing gene annotation resources. Genome Biol 10: R130

Yamagishi Y, Oya K, Matsuura A, Abe H (2018) Use of CK-548 and CK-869 as Arp2/3 complex inhibitors directly suppresses microtubule assembly both in vitro and in vivo. Biochem Biophys Res Commun 496: 834–839

Yang Q, Zhang XF, Pollard TD, Forscher P (2012) Arp2/3 complex-dependent actin networks constrain myosin II function in driving retrograde actin flow. J Cell Biol 197: 939–956

Yoon YJ, Han YM, Choi J, Lee YJ, Yun J, Lee SK, Lee CW, Kang JS, Chi SW, Moon JH et al (2019) Benproperine, an ARPC2 inhibitor, suppresses cancer cell migration and tumor metastasis. Biochem Pharmacol 163: 46–59

Zimmet A, Van Eeuwen T, Boczkowska M, Rebowski G, Murakami K, Dominguez R (2020) Cryo-EM structure of NPF-bound human Arp2/3 complex and activation mechanism. Sci Adv 6: eaaz7651

