## Supplementary figures 1 to 5 for "CK-666 and CK-869 differentially inhibit Arp2/3 iso-complexes"

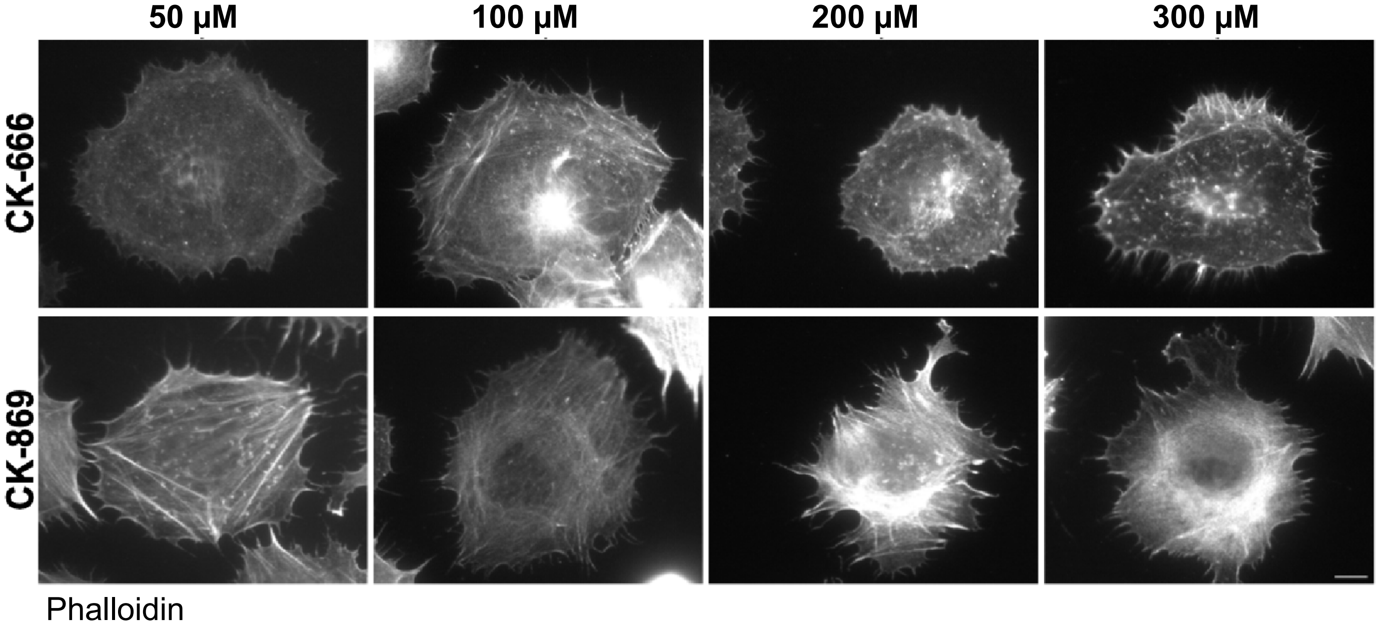


Supplementary Figure 1

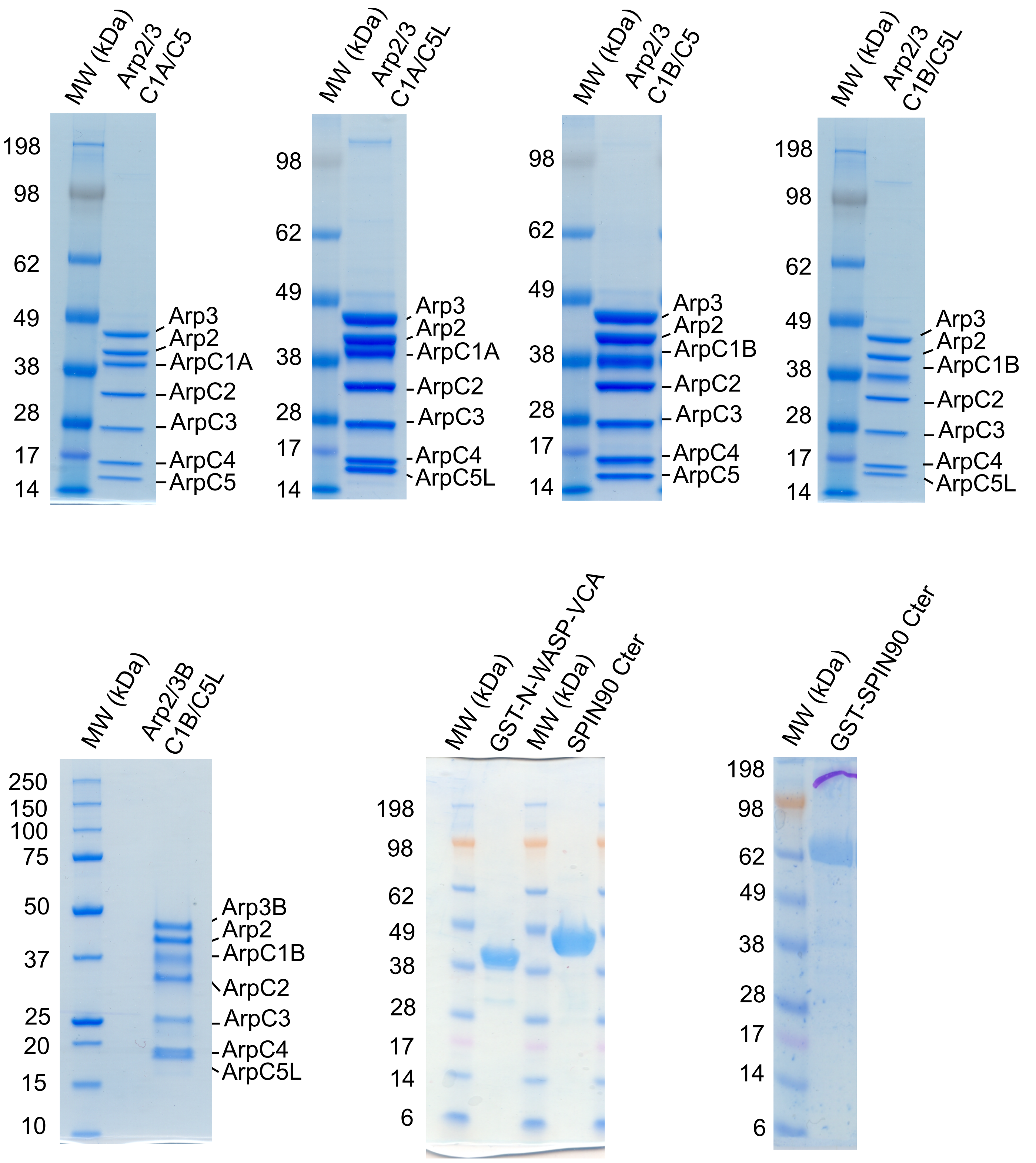


Supplementary Figure 2

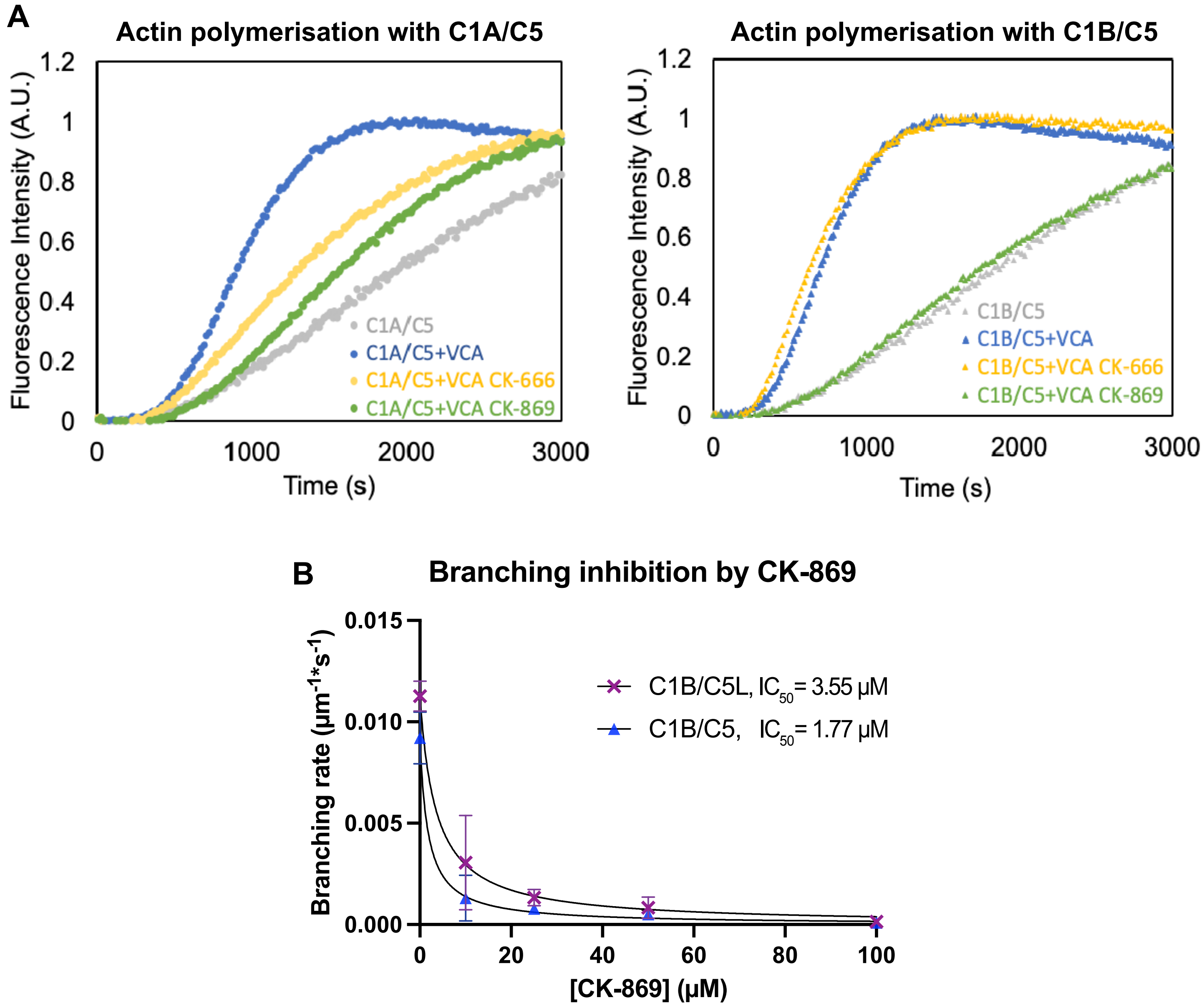


Supplementary Figure 3

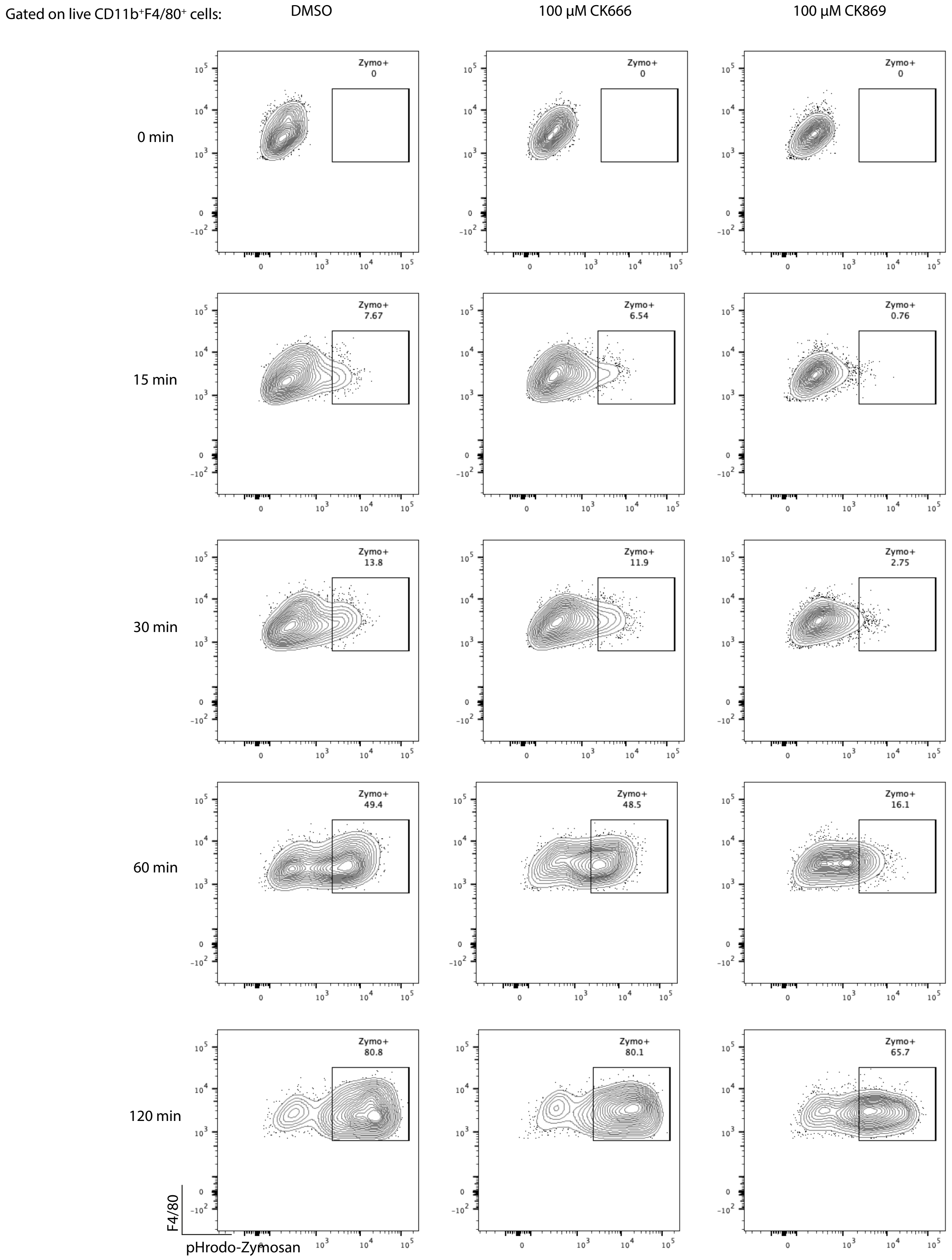


Supplementary Figure 4

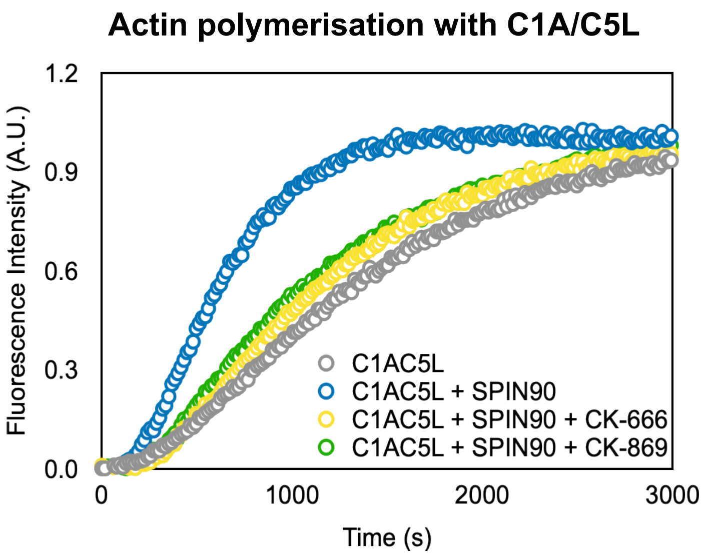


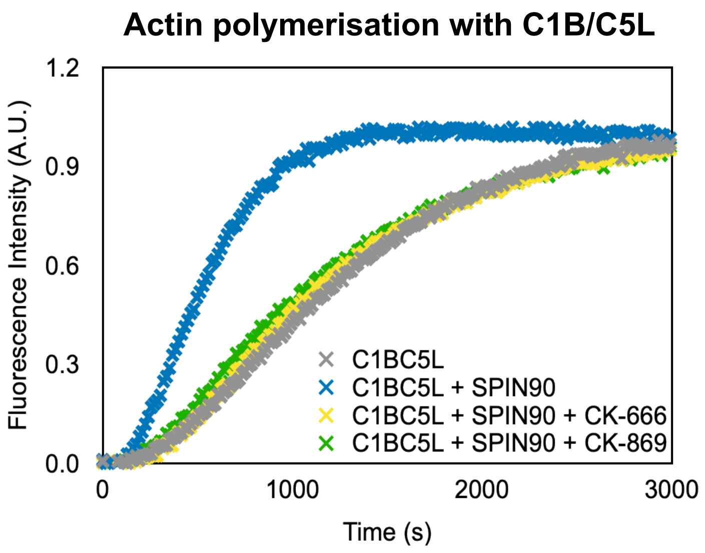


Supplementary Figure 5
